# Effects of spontaneous mutations on survival and reproduction of *Drosophila serrata* infected with Drosophila C virus

**DOI:** 10.1101/2024.04.04.588202

**Authors:** Bonita M. Mendel, Angelique K. Asselin, Karyn N. Johnson, Katrina McGuigan

**Affiliations:** School of the Environment, The University of Queensland, Brisbane QLD 4072, Australia

**Keywords:** LT50, Mutation Accumulation, DsGRP, Wolbachia, immune, productivity

## Abstract

The impact of selection on host immune function genes has been widely documented. However, it remains essentially unknown how mutation influences the quantitative immune traits that selection acts on. Applying a classical mutation accumulation (MA) experimental design in *Drosophila serrata*, we found the mutational variation in susceptibility (median time of death, LT50) to Drosophila C virus (DCV) was of similar magnitude to that reported for intrinsic survival traits. However, the mean LT50 did not change as mutations accumulated, suggestion no directional bias in mutational effects. Maintenance of genetic variance in immune function is hypothesised to be influenced by pleiotropic effects on immunity and other traits that contribute to fitness. To investigate this, we assayed female reproductive output for a subset of MA lines with relatively long or short survival times under DCV infection. Longer survival time tended to be associated with lower reproductive output, suggesting that mutations affecting susceptibility to DCV had pleiotropic effects on investment in reproductive fitness. Further studies are needed to uncover the general patterns of mutational effect on immune responses and other fitness traits, and to determine how selection might typically act on new mutations via their direct and pleiotropic effects.

## Introduction

Pathogens are ubiquitous, with the fitness costs they impose on individual hosts (Price 1980) requiring economically costly interventions (Savary et al. 2019; Antimicrobial Resistance Collaborators 2022) and potentially contributing to the loss of biodiversity (Smith et al. 2006; Preece et al. 2017; Gilbert et al. 2023). Heritable differences in these fitness costs among host individuals has led to the evolution of diverse host defences (Medzhitov et al. 2012; Pereira et al. 2023), and strong signatures of selection on immune function genes (Mukherjee et al. 2014; Shultz and Sackton 2019; Gouy and Excoffier 2020; Larragy et al. 2023). In contrast to this well documented role of selection, we understand little about how mutation contributes to quantitative immune traits, or therefore how the distribution of mutational effects might influence the opportunity for selection and future evolution.

Mutation is the ultimate source of all genetic variation, and the frequency with which mutations arise and their effects on fitness theoretically explain major evolutionary phenomena ranging from sex and recombination (Keightley and Otto 2006) to extinction risk (Lynch and Gabriel 1990). While the influence of mutation on fitness and life history traits has been widely studied (Halligan and Keightley 2009; Conradsen et al. 2022), immune function traits have received little attention. A notable exception is the study by Etienne et al. (2015) on susceptibility (the median lethal time, LT50) of *Caenorhabditis elegans* to a generalist bacterial pathogen, *Pseudomonas aeruginosa*. Mean LT50 did not evolve under mutation and drift, suggesting a similar frequency of mutations increasing and decreasing susceptiblity (Etienne et al. 2015). This contrasts markedly with the general pattern of mutation decreasing fitness (Halligan and Keightley 2009). However, unbiased mutation would provide the fuel for both positive (diversifying) and negative (purifying) selection to act, consistent with evidence from comparative genomic studies that both types of selection have influenced variation in immune function genes (Mukherjee et al. 2014; Early and Clark 2017; Shultz and Sackton 2019; Gouy and Excoffier 2020; Larragy et al. 2023).

Selection acts directly on immune function via pathogen-driven decreases in survival and reproduction. If immune function traits share variation (i.e., are correlated) with other fitness-enhancing traits (e.g., reproduction), selection on those traits can indirectly influence variation in immunity. Comparative genomic studies have provided evidence of this: immune function loci with pleiotropic effects on developmental functions or essential cellular processes evolve slowly relative to loci with immune-limited functions (Enard et al. 2016; Williams et al. 2023).

In general, the competing energetic costs of immune functions and other fitness-enhancing traits are expected to determine the opportunity for selection to shape variation (Viney et al. 2005; Schwenke et al. 2016). At the phenotypic level, immune responses are typically negatively correlated (i.e., trade-off) with changes in other fitness traits, consistent with competing energetic costs (Viney et al. 2005; Schwenke et al. 2016; Albery et al. 2021). How genetic (co)variation among fitness-enhancing traits might reflect energetic demands is expected to be more complex, depending both on how individual traits contribute to fitness and on the genetic architecture (Charlesworth 1990; Houle 1991; Hughes and Leips 2017). There is considerable heterogeneity in reported genetic correlations between immune function and other fitness traits (Ye et al. 2009; van der Most et al. 2011; Faria et al. 2015; Buzatto et al. 2019). Trait correlations due to new mutations, unsorted by selection, may provide insight into the genetic architecture, and how mutations affecting immunity influence direct and indirect selection pressure on the population.

Several studies have linked fitness effects of mutations with immune response by treating infection as a stressor; pathogen infection increases the strength of selection against some, but not all mutations (e.g., Cooper et al. 2005; Young et al. 2009). However, these studies have not considered the variation among host genotypes in their immune response, which is necessary to understand how the interaction between mutation and selection shapes genetic variation in immune responses. In *C. elegans*, Etienne et al. (2015) reported positive mutational correlation between susceptibility (LT50) and fitness (measured in the absence of infection): mutations causing low relative fitness also caused more rapid death under *P. aeruginosa* infection. This suggests that mutations increasing host susceptibility will be under purifying selection via their general fitness effects in the absence of exposure to the pathogen. Notably however, the relationship between fitness and susceptibility was curvilinear, such that longer lifespan of infected hosts was observed across a range of fitness values (Etienne et al. 2015), suggesting some mutations decouple immune and non-immune fitness effects.

In this study, we aimed to contribute to the currently sparse literature on the variation in quantitative immune phenotypes that is introduced by mutation each generation, and how selection both in the presence and absence of the pathogen might influence the persistence of mutations to standing genetic variation. Using a classical mutation accumulation (MA) experiment in *Drosophila serrata*, we estimated the mutational variance in susceptibility (LT50) under systemic infection with Drosophila C virus (DCV). We investigated the correlated mutational effects on fitness by measuring lifetime reproductive output of infected and uninfected females with mutational breeding values associated with relatively high versus low DCV susceptibility.

## Methods

### FLY POPULATIONS

A mutation accumulation (MA) experiment was established from one of the highly inbred *Drosophila serrata* Genome Reference Panel lines (DsGRP: Reddiex et al. 2018). Each of 65 MA lines was maintained by brother-sister mating for 30 generations following protocols detailed in McGuigan et al. (2011). All flies were maintained on a diet of torula yeast and cane sugar (Kannan et al. 2023) at 25°C with a 12-hour light/dark cycle. The intracellular bacteria *Wolbachia* commonly infects insects, and influences host responses to other infections (Hilgenboecker et al. 2008; Hoffmann 2020). We therefore confirmed the experimental population was not infected with *Wolbachia* (Supplementary Information).

Prior to phenotypic data collection, inbreeding was stopped in the MA lines, and all lines (MA and DsGRP-120) were maintained at a census population size of ∼90, in three vials of ∼30 flies (mixed each generation). The rationale for this was twofold. Firstly, it was logistically prohibitive for us to maintain MA inbreeding while conducting the phenotype assays. Secondly, the increased density imposed a resource limitation (McGuigan 2009). Low larval density and benign laboratory conditions, limiting (or eliminating) resource competition, may result in underestimation of mutational effects (Agrawal and Whitlock 2010; Kannan et al. 2023).

The increased population size of each line will alter the mutation-selection-drift dynamics. Notably, populations as small as *N* = 10 have been empirically shown to halt deleterious mutation fixation (Estes et al. 2004; Katju et al. 2015; Luijckx et al. 2018). While beneficial mutations may, in contrast, rapidly sweep to fixation when *N* > 2, such mutations appear to be relatively rare (Keightley and Lynch 2003; Halligan and Keightley 2009) and sampled infrequently even in large populations (*N* > 1000) (e.g., Denver et al. 2010). Thus, with *N* ∼ 90, we expect among MA line variation will predominantly reflect mutations accumulated during the 30 generations of brother-sister mating, while mutations arising after population size expansion will contribute to within-line variance. In estimating mutational variance, we accounted for the larger census population size as detailed below.

### EXPERIMENT 1: DCV INFECTION SUCEPTIBILITY

Drosophila C virus (DCV) is a well-studied virus that naturally infects members of the *Drosophila* genus (Kapun et al. 2010). DCV infected *D. melanogaster* have been collected (Christian 1992) from within the natural range of *D. serrata* (Schiffer et al. 2004). However, DCV infection of wild or laboratory *D. serrata* has not been reported. We therefore first confirmed that DCV had pathogenic effects on *D. serrata*.

Abdominal injection is a commonly used to study DCV infection, ensuring a standard dose is delivered to each individual fly (Osborne et al. 2009). In the current study, individual 5-7 day old virgin flies were abdominally injected with 56 nl of DCV (1 × 10^8^ infectious units per millilitre) with a Nanoject II microinjector (Drummond) using a pulled glass capillary (Hedges and Johnson 2008; Osborne et al. 2009). Due to logistical limits on sample size, we focus our investigation on only one sex, females. While we confirmed males experience similar mortality following DCV infection (Supplementary Material) we note that mutational effects may differ between the sexes. We focus on females due to their potentially greater influence on population dynamics under infection. Specifically, evidence from *D. melanogaster* suggests females are both less able to clear DCV infection and more likely to transmit it to other adult flies than males are (Gomariz-Zilber et al. 1995). Further, female fecundity may have a greater impact on population growth rates than male reproduction does (Whitlock and Agrawal 2009).

To determine whether *D. serrata* was a susceptible host of DCV, we randomly sampled 15 MA lines and exposed them to DCV or phosphate-buffered saline, PBS (a control for injection-related mortality). For each line, four replicate vials of seven virgin females were randomly assigned to an infection treatment (DCV or PBS). Injections were conducted over two sequential days, with one replicate per infection treatment per day. Flies were transferred to fresh media within 24 hrs of injection, and again after six days to ensure deteriorating conditions did not contribute to mortality. Mortality observed on the day following injection (< 10% of the 420 flies injected) was assumed to be due to factors other than infection; these flies were excluded from the initial (day 1) census of the vial, and thus from all analyses. Vials were monitored daily, and the number of days post-injection (dpi) each individual fly survived was recorded. Data collection ended when all DCV-injected flies had died (11 dpi); surviving PBS flies were censored on this day.

The null hypothesis that PBS and DCV injected flies experienced the same rate of mortality was tested using the coxme package (Therneau 2022) in R (R Core Team 2022) using RStudio (Posit Team 2023) to fit the mixed effects model:

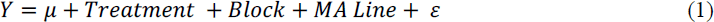

where *Y* was a vector of the day on which each individual died or was censored, *μ* the global mean survival; treatment (PBS or DCV injected) and block (1^st^ or 2^nd^ day of injection) were both modelled as categorical fixed effects, while MA line and replicate vial (nested within MA line and treatment, ϵ) were modelled as random effects. We applied a log-likelihood ratio test (LRT), comparing the fit (-2 log likelihood) of model (1) to a nested model in which injection treatment was not fit; the difference in fit between nested models follows a chi-square distribution with degrees of freedom equal to the difference in number of model parameters (here, 1) (Self and Liang 1987). The survminer R package (Kassambara et al. 2021) was used illustrate the rate of mortality in each treatment from a proportional hazards model.

### EXPERIMENT 2: MUTATIONAL VARIANCE IN DCV INDUCED MORTALITY

Having confirmed *D. serrata* susceptibility to DCV (see Results), we investigated how mutation affected survival time following DCV infection. Four replicate vials (seven virgin females per vial) per MA line, and twelve replicate vials for the DsGRP-120 ancestor were injected with DCV following the protocols detailed above. Injections were conducted over four sequential days; one replicate vial per MA (and three for DsGRP-120) were injected each day. The 68 vials (65 MA lines plus three replicates of DsGRP-120) were randomly ordered on each injection day. A total of 1,904 flies in 272 vials were injected. Survival of all flies within each vial was recorded daily as detailed above. Flies were transferred to fresh media every four days.

We quantified the effect of DCV infection on survival for each of the 272 vials as the median lethal time (LT50: time in days to mortality of 50% of the seven females per vial). To estimate LT50, a generalised linear model with a binomial error distribution and a logit link function (implemented using the glm function in base R: R Core Team 2022) was fit to the data for each vial separately (i.e., 272 models):

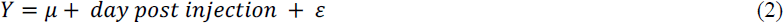

where *Y* was a binomial response vector of number of flies (dead or alive) in that vial each day, *μ* (*β* _0_) was the intercept, day post injection was a continuous fixed effect predictor, and ε was the unexplained residual. The dose.p function in the MASS package (Venables and Ripley 2002) was used to calculate LT50 for each vial, which under the logit link function simplifies to LT50 = −*β* _0_/*β* _1_, where *β* _0_ and *β* _1_ are the model estimates of the intercept and slope, respectively. For 14 vials (each from a different MA line), death was precipitous such that all individuals died on the same, or consecutive days. This data distribution resulted in convergence problems, and unreliable estimates of LT50. These vials were excluded from all analyses. Across the remaining 258 vials, LT50 estimates followed a Gaussian distribution. These LT50 data were analysed to determine the effect of mutation on DCV infection survival time i) variance and ii) population mean.

To estimate the among-line (mutational) variance in LT50, we excluded DsGRP-120, and used PROC MIXED in SAS (SAS Institute Inc. 2012), with restricted maximum likelihood (REML), to fit:

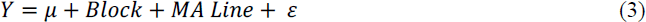

where *Y* was the vector of LT50 values per vial, *μ* was the global mean LT50, block was a categorical fixed effects capturing the effect of the sequential injection days (1 through 4), MA Line was fit as a random effect to estimate the among-line variance (*V*_L_) and ϵ was the unexplained residual variance among replicate vials per MA line. To test the hypothesis that *V*_L_ was distinct from zero (i.e., that mutations affecting DCV susceptibility had accumulated), a log-likelihood ratio test (LRT) was applied, comparing model (3) to a model in which *V*_L_ was constrained to zero. As variances cannot be negative, this hypothesis test is one tailed; the test statistic follows a chi-square distribution with a mixture of 0,1 degrees of freedom (Self and Liang 1987). To place robust confidence intervals on *V*_L_ and derived parameters (detailed below), we adapted the REML-MVN approach developed by Meyer and Houle (2013); (Houle and Meyer 2015). Using the rnorm function in base R (R Core Team 2022), we obtained 10,000 samples from the normal distribution *N(μ,*σ^2^) where *μ* was defined as the REML estimate of V_L_ (or *V*_E_, the residual variance) and σ^2^ as the corresponding inverse Fisher information parameter, retained from the REML model fit.

We estimated the per-generation rate of increase of phenotypic variance due to mutation (i.e., the mutational variance) as *V*_M_ = *V*_L_/2*t*, where *V*_L_ is the among-line variance from model (3), and *t* is the number of generations of mutation accumulation. This relationship defines *V*_M_ under brother-sister mating (Lynch and Walsh 1998), where the small effective population size ensures rapid fixation (or loss) of mutations. However, survival assays were conducted after seven generations of relaxed inbreeding (each line expanded to ∼90 flies). The effective population size was unknown during these seven generations, precluding accurate calibration of the expected rate of mutation accumulation. We aimed to nonetheless gain some insight into the magnitude of *V_M_* relative to published estimates by setting *t* = 30 (generations of brother-sister mating) and *t* = 37. Assuming that the increased population size was effective at preventing any further mutations fixing after population expansion (i.e., *t* = 30) defines the maximum *V*_M_, while assuming mutations accumulated at the same rate over the 30 generations of brother-sister mating and the seven generations of relaxed inbreeding (*t* = 37) defines the minimum *V*_M_.

We placed these estimates of *V_M_* on dimensionless scales for comparison by estimating the mutational heritability (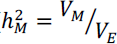, where *V_E_*, the environmental variance, was the residual, among replicate vial, variance), coefficient of mutational variance (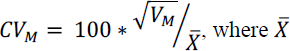 was the mean LT50 of MA flies), and a second mean-standardised metric, opportunity for selection 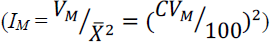 (Houle 1992; Garcia-Gonzalez et al. 2012). Confidence intervals (CI) were calculated for 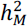 using the REML-MVN samples described above, obtained for both *V_L_* and *V_E_* (residual variance). To obtain CI of mean standardized parameters, we used the rnorm function to draw 10,000 samples from the normal distribution defined by the mean and standard deviation of LT50 of the MA lines. While we report the mutational heritability of LT50, we note that, because LT50 was estimated from information on seven females per vial, we likely under-estimate the environmental variance in susceptibility, and this downward bias in the estimate of the denominator upwardly biases the estimate of 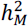.

Next, we determined whether mutations affected the population mean LT50. If the frequency or average effect size of mutations were biased toward either lower or higher trait values, the trait mean of the MA lines would diverge from that of their ancestor. To test the null hypothesis that mean LT50 of the 65 MA lines was the same as mean LT50 of the ancestor, DsGRP-120, we followed Latimer et al. (2014) in applying a bootstrapping approach to accommodate the unbalanced data (here, 1 ancestor, 12 replicate vials versus 65 MA lines, 258 replicate vials). We generated 1,000 datasets by randomly sampling (with replacement) three MA line and three DsGRP-120 vials per injection day (24 vials in total), using PROC SURVEYSELECT in SAS (SAS Institute Inc. 2012). We then used maximum likelihood (ML), implemented using PROC MIXED in SAS, to fit:

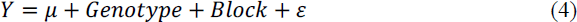

where *Y* was the vector of LT50 values for each vial, *μ* and block were as described for model (3), while genotype was a categorical fixed effect capturing the effect of MA panel versus DsGRP-120 ancestor. The residual, ϵ, again modelled unexplained variation among vials; MA line was not accounted for as random sampling of 12 vials rarely resulted in replicate measures per MA line. This model was fit 1,000 times, to each random sample, and the difference between least squares means (ι1M = Ancestor LSMean - MA LSMean) retained. We inferred trait evolution under mutation where 95% of bootstrap estimates did not include zero (no difference).

### EXPERIMENT 3: DCV RESISTANCE AND PRODUCTIVITY TRADE-OFF

Finally, we investigated whether mutations affecting susceptibility to DCV (i.e., LT50) might have pleiotropic effects on reproductive fitness. We investigated these potential costs of immune responses in both the presence and absence of DCV infection. Twelve MA lines were selected based on their LT50 (estimated as detailed above): six with high LT50 values (median = 7.82), reflecting relatively low DCV susceptibility, and six with low LT50 (median = 4.79), reflecting relatively high susceptibility. For these 12 MA lines, and their ancestor DsGRP-120, a total of 12 replicate vials with three virgin females were collected (156 vials total).

Six vials per line were assigned to a PBS treatment and six to a DCV treatment, and flies injected following the protocols detailed above. The day after injection, females were moved to new vials, with two virgin males from the same line. Each day, flies were moved to a new vial to allow us to determine the temporal distribution of reproductive output. We limited the potential contribution of male harassment to female mortality by removing one (randomly chosen) male when female mortality resulted in a male-biased sex ratio (i.e., one surviving female plus two males). We terminated the experiment at 12 days post injection (dpi), when females were 16 days old, and expected to have reached reproductive senescence.

We counted the number of adults eclosing in each vial (i.e., for each day of laying of each injection replicate per line). This measure of productivity captures effects of both female fecundity and embryo survival to adulthood. We observed very little mortality at the pupal stage (number of eclosed adults was very similar to the number of pupae: K. McGuigan, unpublished data), but could not further disentangle fecundity and survival effects within this experiment. For each vial, we accounted for death of females (i.e., reduction in the number of females potentially contributing offspring) by dividing the count of eclosed offspring by the number of potential mothers on that day.

The productivity metrics were not normally distributed, and we thus applied non-parametric analyses. First, paired Wilcoxon rank sum tests were applied in R (R Core Team 2022; Posit Team 2023) to test the hypotheses that DCV infection shortened the reproductive period (last day on which offspring were produced as the response variable) or decreased the number of offspring (the maximum number of offspring per female on any day as the response variable). The 13 lines (12 MA plus ancestor) were the paired observations, and the analyses were applied to the median value of the six replicate vials per line per injection treatment.

Next, we tested our focal hypothesis that lines classified as having high DCV susceptibility (low LT50 values in Experiment 2) produced a different number of offspring than the lines classified as having low DCV susceptibility (high LT50 values in Experiment 2). The ancestor line was excluded from these analyses (but retained for plotting). Separately for PBS and DCV injected flies, we applied unpaired Wilcoxon rank sum tests to the maximum number of offspring per female on any day, and to the lifetime reproductive output of each female.

## Results

### SUSCEPTIBLE TO DCV AND MUTATIONAL VARIATION IN SURVIVAL TIME OF INFECTED FLIES

Experiment 1, in which flies from a random sample of 15 MA lines were exposed to either PBS (injection control) or live virus, confirmed *D. serrata* is a susceptible host of DCV. Injection with DCV significantly increased the rate of mortality relative to injection with PBS (LRT of Treatment: χ^2^ = 358.92, d.f. =1, *P* < 0.0001; hazard ratio = 3.31 ± 0.22; Figure 1). All DCV-infected flies died within 11 days of being injected, while only 23% mortality was observed for the control, PBS-injected, flies within the same period (Figure 1).

**Figure 1.**
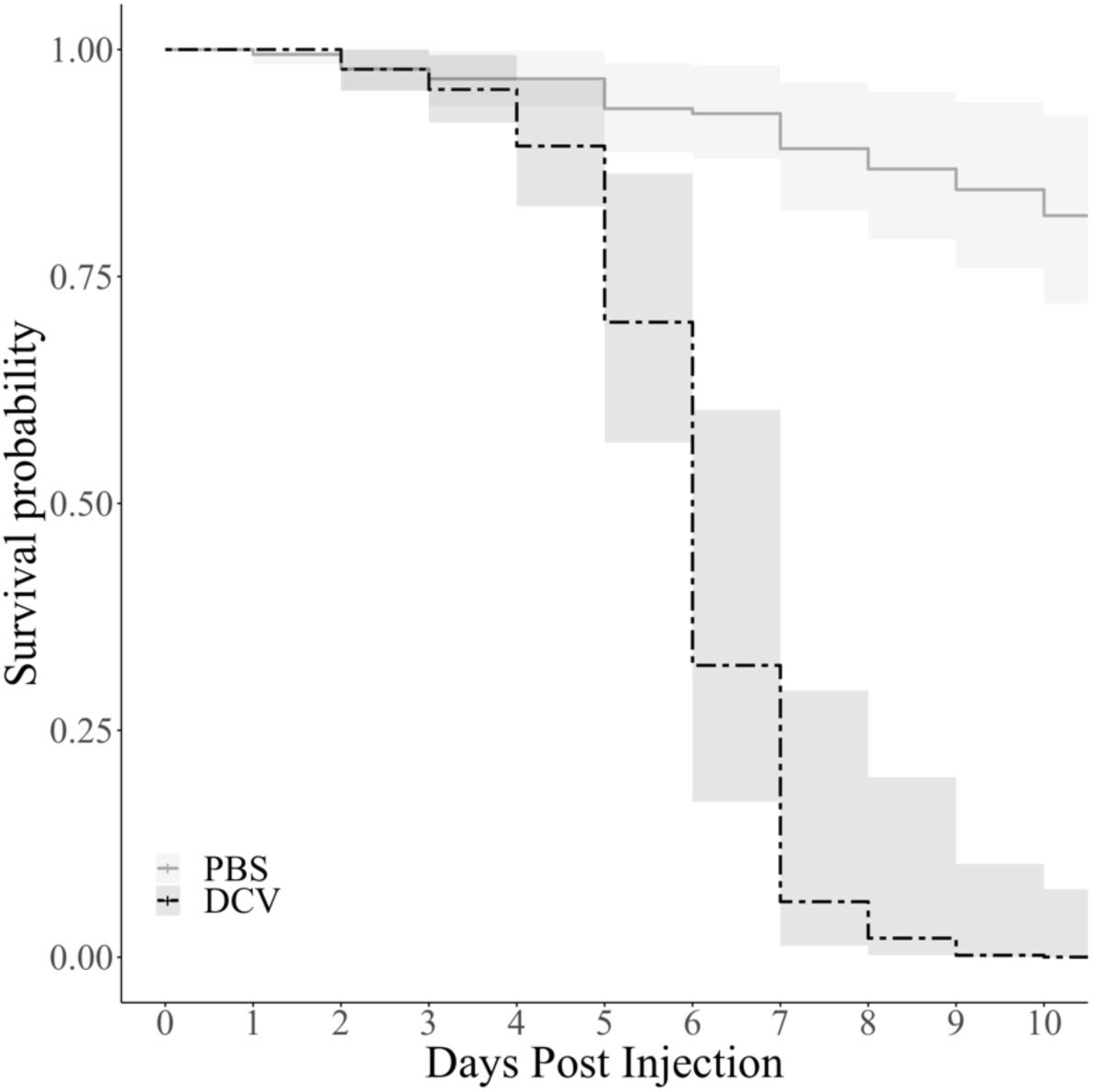
Survival of *D. serrata* challenged with PBS or DCV. For females from 15 randomly chosen MA lines, the probability of being alive each day following injection with the pathogen, DCV (dashed black line), or a placebo (PBS; solid grey line) was estimated from a Cox’s proportional hazard model. Shading indicates 95% CI. All DCV injected flies were dead by day 11, and PBS flies were censored.

There was strong statistical support for mutational (among-line) variance in DCV susceptibility (Experiment 2, LRT of zero among-line variance: χ^2^ = 14.30, d.f. = (0,1), *P* < 0.0001) (Table 1; Figure 2A). Across all 65 MA lines, the average LT50 (time to 50% mortality of seven females in a vial following DCV injection) was 6.29 days (Best Linear Unbiased Predictors, BLUPs, ranged from 4.96 to 7.32 days) (Figure 2A). While LT50 of the ancestor DsGRP-120 was slightly longer (6.40 days), we accepted the null hypothesis that Λ1M = (Ancestor - MA) = 0 (*P* = 0.6440; Figure 2B), and that there was no bias in the direction of mutational effects.

**Figure 2.**
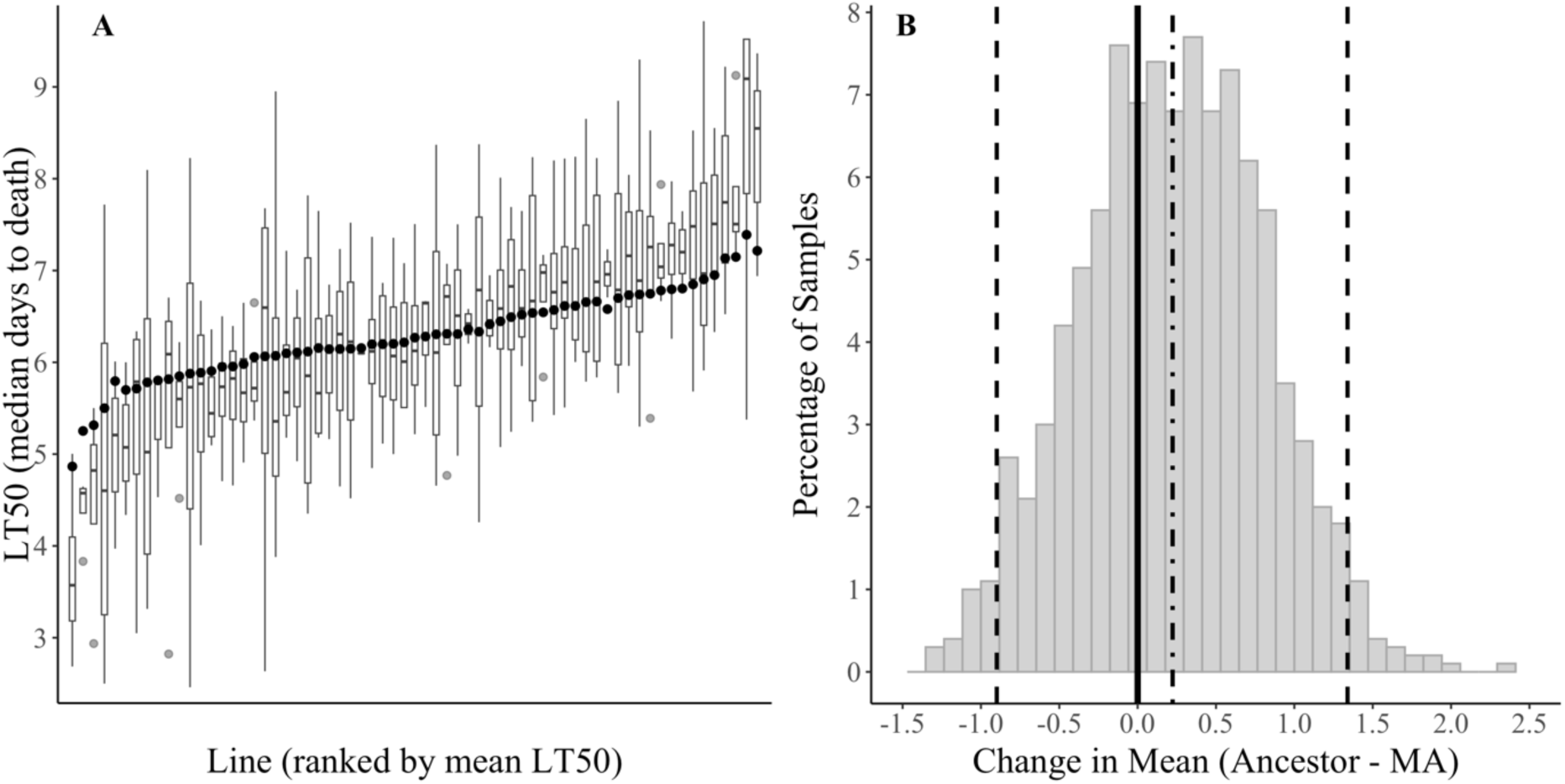
Distribution of LT50 values of MA lines and DsGRP-120. A) The 65 MA lines were ranked by their mean LT50, and their Best Linear Unbiased Predictors (BLUPs from model 3; solid black circles) plotted over boxplots of the LT50 distribution of the four replicate vials assayed per line. Boxplots indicate the median (black bar), interquartile range (IQR) (box), 1.5IQR (whiskers) and extreme values (grey shaded circles). B) The distribution of the difference in mean LT50 between the ancestor and MA population (DsGRP-120 minus MA) of 1,000 stratified random samples. The vertical lines indicate zero (solid line; represents the null hypothesis of no directional bias of mutational effects), upper and lower 95% percentiles (dotted line), and the median (dashed line).

**Table 1.**
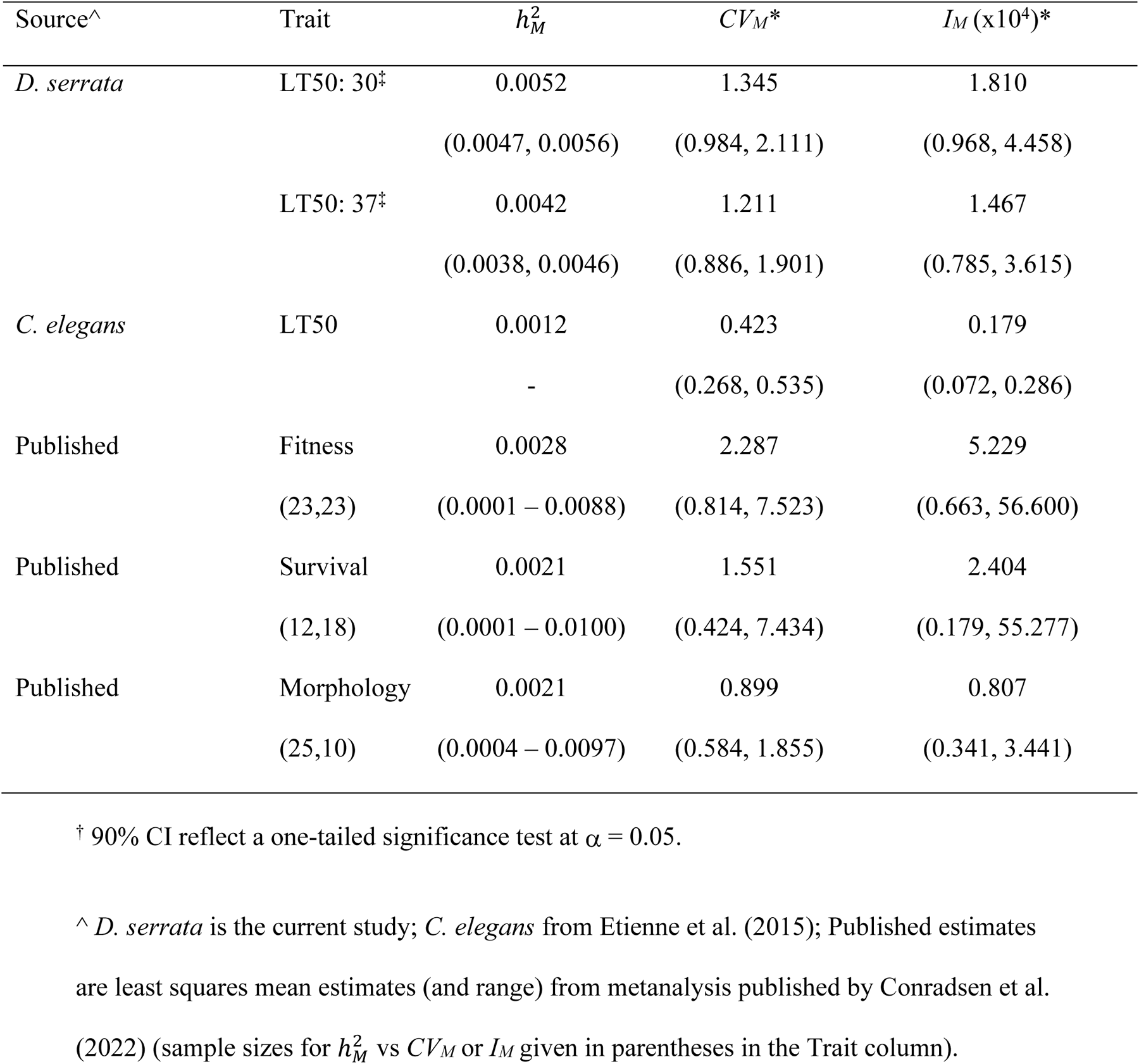
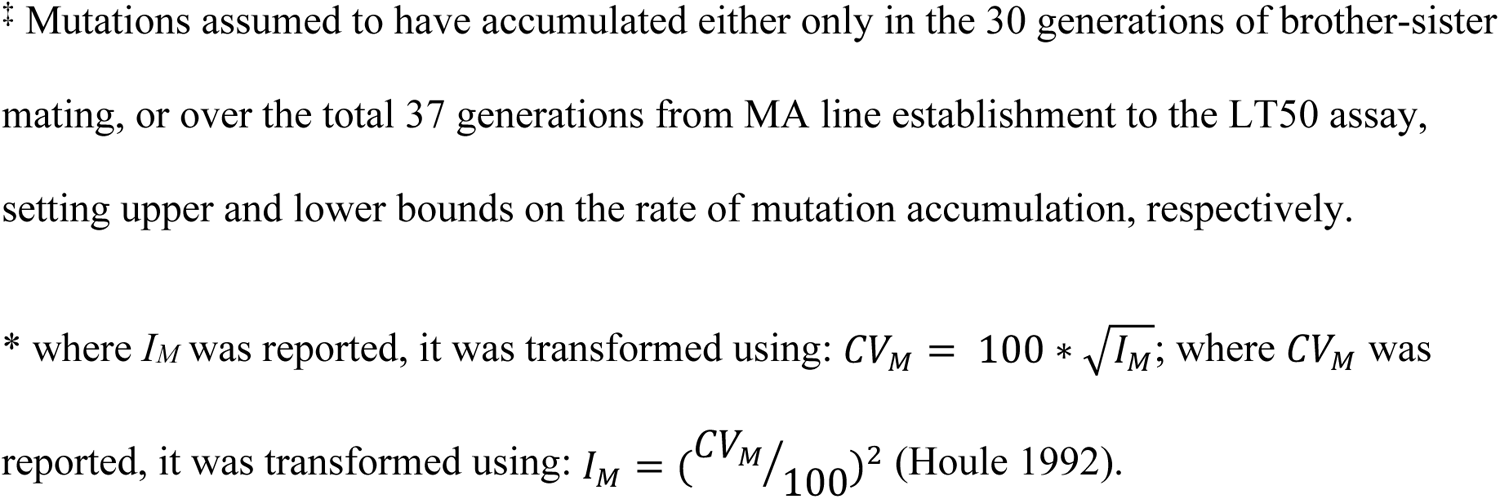
Estimates of scaled mutational variance. Values in parentheses below estimates depend on source: REML-MVN 90% CI^†^ for this study (*D. serrata* LT50 under DCV infection), 90% CI from reported standard error for *C. elegans* (not available for 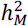), and the minimum and maximum across published estimates.

| Source<sup>^</sup> | Trait | $h_M^2$ | $CV_M^*$ | $I_M$ (x10<sup>4</sup>)* |
| --- | --- | --- | --- | --- |
| *D. serrata* | LT50: 30<sup>‡</sup> | 0.0052 | 1.345 | 1.810 |
|  |  | (0.0047, 0.0056) | (0.984, 2.111) | (0.968, 4.458) |
|  | LT50: 37<sup>‡</sup> | 0.0042 | 1.211 | 1.467 |
|  |  | (0.0038, 0.0046) | (0.886, 1.901) | (0.785, 3.615) |
| *C. elegans* | LT50 | 0.0012 | 0.423 | 0.179 |
|  |  | - | (0.268, 0.535) | (0.072, 0.286) |
| Published | Fitness | 0.0028 | 2.287 | 5.229 |
|  | (23,23) | (0.0001 – 0.0088) | (0.814, 7.523) | (0.663, 56.600) |
| Published | Survival | 0.0021 | 1.551 | 2.404 |
|  | (12,18) | (0.0001 – 0.0100) | (0.424, 7.434) | (0.179, 55.277) |
| Published | Morphology | 0.0021 | 0.899 | 0.807 |
|  | (25,10) | (0.0004 – 0.0097) | (0.584, 1.855) | (0.341, 3.441) |

<sup>†</sup> 90% CI reflect a one-tailed significance test at $\alpha = 0.05$ .
<sup>^</sup> *D. serrata* is the current study; *C. elegans* from Etienne et al. (2015); Published estimates are least squares mean estimates (and range) from metanalysis published by Conradsen et al. (2022) (sample sizes for $h_M^2$ vs $CV_M$ or $I_M$ given in parentheses in the Trait column).
‡ Mutations assumed to have accumulated either only in the 30 generations of brother-sister mating, or over the total 37 generations from MA line establishment to the LT50 assay, setting upper and lower bounds on the rate of mutation accumulation, respectively.
\* where $I_M$ was reported, it was transformed using: $CV_M = 100 * \sqrt{I_M}$ ; where $CV_M$ was reported, it was transformed using: $I_M = (CV_M/100)^2$ (Houle 1992).

### EFFECT OF DCV ON FEMALE PRODUCTIVITY (EXPERIMENT 3)

DCV infection very strongly impacted female reproductive output by truncating the reproductive period. No DCV injected females produced any offspring beyond 6 days post injection (dpi), while some PBS injected females were still producing offspring at 11 dpi. The DCV treatment truncated the median reproductive period by 4.5 days (Figure 3A; Wilcoxon *W* = 0.0, *P* = 0.0058, *N* =13; note: the test statistic value of zero indicates that, for all 13 assayed lines, reproductive period was shorter for DCV than PBS). As DCV reduces lifespan (Figure 1), we further investigated whether the observed difference in reproductive lifespan was a simple consequence of earlier mortality of DCV infected flies. Considering only vials with at least one offspring (*N =* 92), we determined the last day of reproduction as a proportion of lifespan (maximum 12 dpi due to censoring). While DCV injected flies ceased reproduction at 22% of their lifespan, PBS injected flies, on average, bred until 52% of their lifespan, a significant difference in reproductive span (*W* = 7.0, *P* = 0.0046, *N* =13).

**Figure 3.**
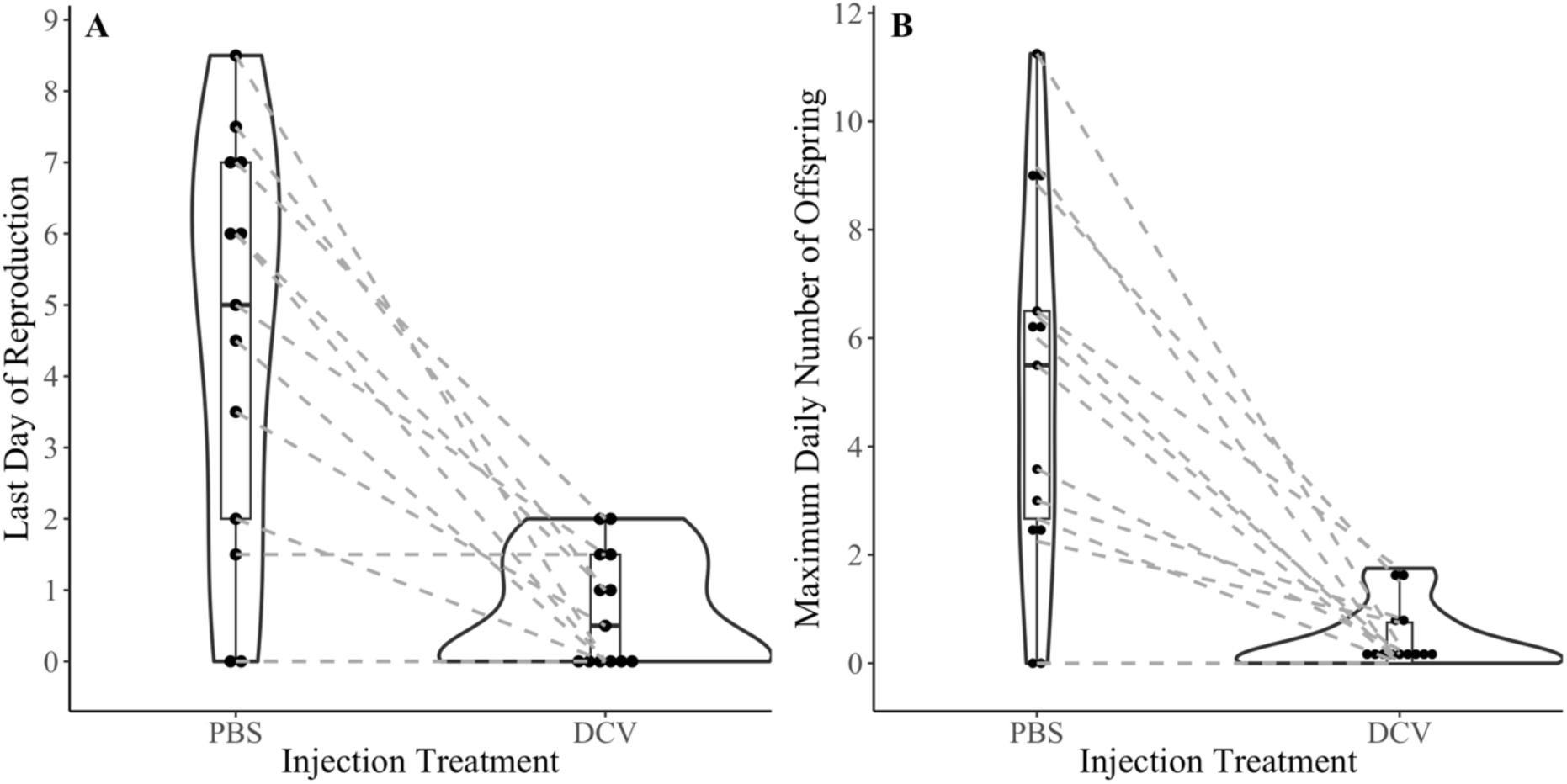
Effect of virus treatment on female productivity. A) Duration of the reproductive period, indexed as the last day on which females produced offspring. B) The maximum number of offspring produced by a female in a 24hr period. Solid black points are the median value (of six replicate vials) of each of the 13 lines assayed. Dotted lines connect the observations of a line under each injection treatment. The violin plots illustrate data density, and are overlain on boxplots (box is the interquartile range, IQR; horizontal bar the median over the 13 lines; whiskers 1.5 IQR).

DCV infection also reduced the maximum number of offspring a female produced in a day. Most (43 of 78) vials of DCV injected females produced no offspring, while reproductive failure was half of this for the PBS treatment (21 of 78 initiated vials). Reflecting the high reproductive failure under DCV infection, the maximum number of offspring females produced per day was significantly lower under DCV than PBS (median = 0.17 and 5.50, respectively; Wilcoxon *W* = 0.0, *P* = 0.0039, *N* =13; Figure 3B).

For uninfected flies (PBS injected), we accepted the null hypotheses that there was no difference between females from high (low LT50) and low (high LT50) susceptibility lines in their reproductive lifespan (Figure 4A; Wilcoxon *U* = 13, *P* = 0.4688). Similarly, our data did not support a difference in maximum daily reproductive output per female (Figure 4C; Wilcoxon *U* = 14.5, *P* = 0.6304). Under DCV infection, there was weak statistical support for higher reproductive output of high susceptibility lines (last day of reproduction: Wilcoxon *U* = 6.5, *P* = 0.0590, Figure 4B; maximum daily reproductive output per female: Wilcoxon *U* = 6.5, *P* = 0.0601, Figure 4D). This result reflects higher overall rate of complete reproductive failure of low susceptibility lines: five of the six lines had a median of zero for the maximum daily reproductive output per female (Figure 4B). Indeed, of the 36 DCV-injected vials from the six low susceptibility MA lines, most (24, 67%) had no offspring; in contrast, 17 (47%) of vials from high susceptibility MA lines did not produce any offspring. The ancestor line, DsRGP-120, was not included in these analyses (as there is only one line), but had noticeably higher reproductive success (only 25% of vials failed to produce any offspring), and higher median daily output than most lines under both control (PBS: Figure 4A), and infected (DCV: Figure 4B) conditions, consistent with mutations typically having deleterious effects on fitness.

**Figure 4.**
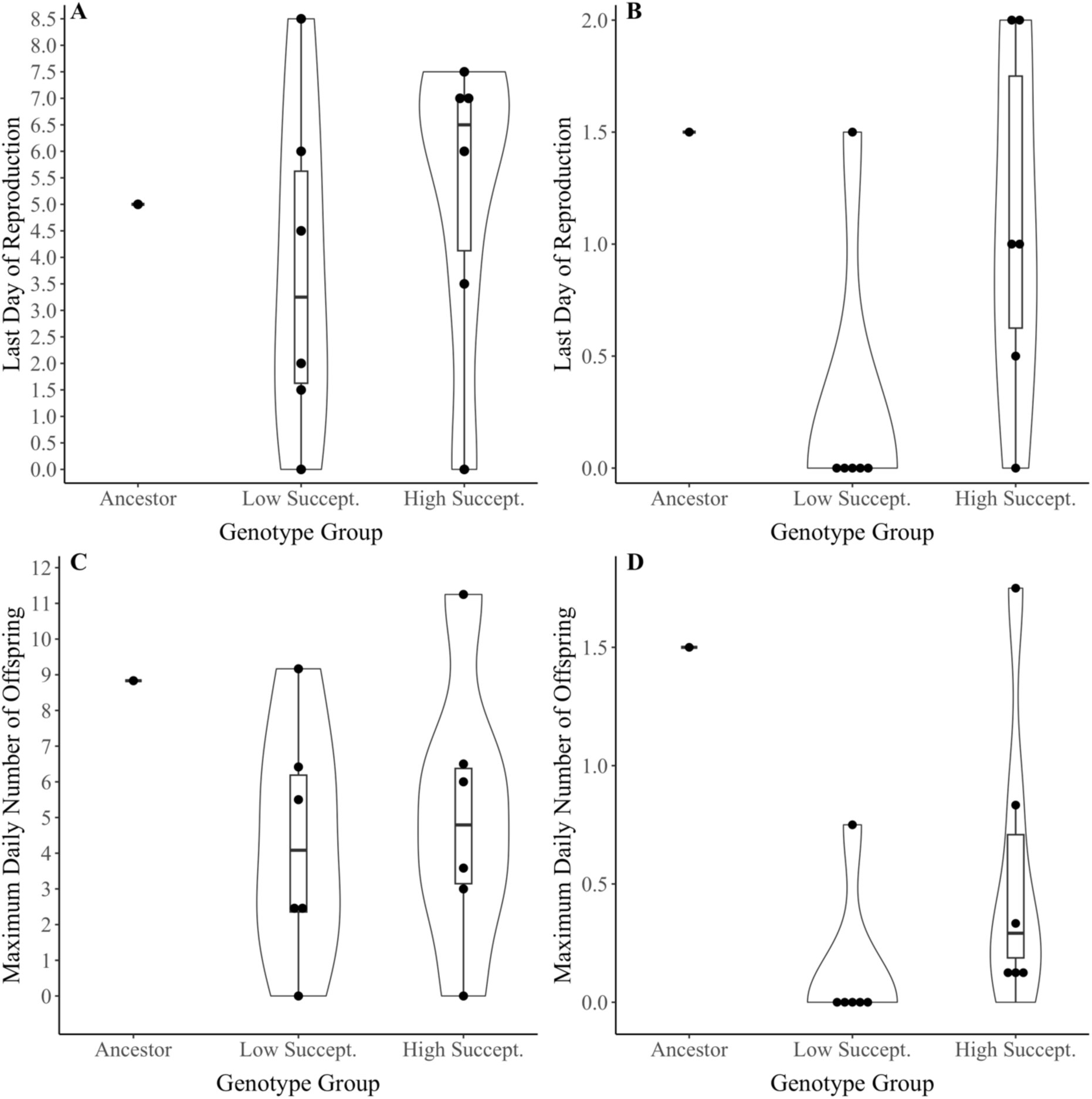
Productivity of high and low DCV susceptibility lines and their ancestor. For (A,C) PBS injected flies and (B,D) DCV injected flies, the line medians of (A,B) the maximum reproductive lifespan (last day of reproduction) or (C,D) the maximum number of offspring per female per day are plotted. Points, violin and boxplots are as described for Figure 3. Note: the ancestor is plotted, but was not considered in statistical analyses.

## DISCUSSION

Pathogen infection has fitness costs to their host, as evidenced by evolution of host defences (Kimbrell and Beutler 2001). Despite these fitness consequences, relatively little attention has been paid to the nature of mutational input to quantitative immune function traits. Considering spontaneously arising mutations in the (near) absence of selection in *D. serrata*, we found: i) mutation introduced variation for median survival time (LT50) following exposure to DCV at a rate similar to that reported for other trait types; ii) mutations had unbiased effects on LT50 and; iii) mutations impacting susceptibility to DCV may have pleiotropic effects on reproductive fitness. We consider these results in detail below.

The mutational variance for *D. serrata* DCV LT50 was comparable to mean-standardised (*CV*_M_ or *I_M_*) published estimates, particularly for intrinsic survival traits (to age, without exposure to pathogens) (Table 1; Conradsen et al. 2022). Etienne et al. (2015) similarly observed LT50 *I_M_* to be only slightly below intrinsic survival measured for the same set of MA lines, but substantially below *I_M_* for total fitness. Differences among traits in the magnitude of mutational variance is predicted to reflect differences in how much of the genome affects them, and thus their mutation rate (Houle 1998). Thus, our results and those of Etienne et al. (2015) suggest that the assayed immune function trait of LT50 may be influenced by less of the genome than total fitness, but a similar portion to that influencing intrinsic survival, and more than influencing morphological traits.

Notably, our estimate of LT50 *I_M_* was more than an order of magnitude above that reported for *C. elegans* LT50 (Etienne et al. 2015) (Table 1). Across a range of different traits, mutational variance is typically substantially lower for nematodes (where data predominantly comes from *C. elegans*) than for *Drosophila* (where data predominantly comes from *D. melanogaster*) (Conradsen et al. 2022). Thus, the difference for LT50 likely reflects a general difference in mutational characteristics between the species. We note, however, that while our investigation focused on a relatively specialised pathogen (Drosophila C virus), Etienne et al. (2015) investigated susceptibility to a generalist pathogen (*Pseudomonas aeruginosa*) that infects a wide taxonomic range of hosts. Investigations of standing genetic variation have suggested greater genetic variance and/or larger effect loci for host susceptibility to specialist or coevolved pathogens (Magwire et al. 2012; Palmer et al. 2018; Duxbury et al. 2019). This difference is likely to reflect the action of selection: an architecture of few, large effect, variants is predicted to evolve under local adaptation (Yeaman 2022). However, future studies of host immune function should assess whether the general characteristics of mutational input (i.e., magnitude of variance; average direction of effect) varies among pathogens, including between taxonomic classes of pathogen (e.g., virus versus bacteria) as well as between pathogens with different co-evolutionary histories with the host.

Consistent with Etienne et al. (2015) we did not detect evidence of mutation bias on immune responses: fixation of spontaneous mutations did not change the population mean susceptibility to DCV systemic infection (Figure 2B; Figure S3). On one hand, this observation is consistent with evidence of both strong positive and negative selection on immune function loci (Mukherjee et al. 2014; Early and Clark 2017; Shultz and Sackton 2019; Gouy and Excoffier 2020; Larragy et al. 2023). On the other hand, similar LT50 of MA and their ancestor was surprising given the high fitness cost of infection (Figure 3), and the pervasive evidence that fitness (survival and reproduction) declines as mutations accumulate (Halligan and Keightley 2009). Our experiment was not designed to test for divergence of MA and ancestor fitness, but we note that reproductive output of DsGRP-120 was greater than most of the 12 sampled MA lines (Figure 4), consistent with the typical pattern of deleterious mutation. Differences in bias in direction of effect of mutations between immune function and fitness (or other fitness-enhancing traits) may arise even where mutations have pleiotropic effects on both traits, although this joint distribution of effects remains poorly characterised for any set of correlated traits (McGuigan and Blows 2013). We note that our MA experiment was of relatively short duration (30 generations), and that relatively small differences in frequency or effect size of mutations increasing versus decreasing susceptibility may become apparent over longer time frames. However, Etienne et al. (2015) found no change in LT50 after 250 generations of mutation.

The evolutionary fate of mutations will depend on their effect on total fitness. While universally true, this is particularly important to consider for immune response traits, which are only expressed, and thus subject to direct selection, in the presence of a pathogen, and where costs of expression will influence both direct and indirect selection. Here, in the absence of the pathogen, we observed no difference in reproductive output between MA lines with relatively high versus low susceptibility to DCV (Figure 4A, C). This result contrasts with the positive correlation between fitness and LT50 reported by Etienne et al. (2015). It also contrasts with the pervasively positive mutational correlations reported among different fitness-enhancing life history traits (Houle et al. 1994; Estes et al. 2005; McGuigan et al. 2011). Positive mutational correlation among life history traits is predicted if mutations predominantly influence ability to acquire and efficiently utilize resources, resulting in similar effects across energetically expensive traits (Houle 1991). Such mutations may be effectively targeted by selection, leading to standing genetic variation composed of mutations with either trait-limited or antagonistically pleiotropic alleles, even if mutations giving rise to them are relatively rarer or of smaller effect size.

Our data did, however, suggest a difference between high and low susceptibility lines in the impact of DCV infection on reproductive output. MA lines with relatively low susceptibility to DCV (i.e., high LT50) had lower reproductive success then high susceptibility lines when infected with DCV (Figure 4B, D). This result may reflect differences in strategic investment of energy resources in mounting an immune response versus reproduction (i.e., mutational effects on loci affecting allocation). High perceived risk of extrinsic mortality (including due to disease) is predicted to select for higher investment in current reproduction (Hirshfield and Tinkle 1975; Corbel and Carazo 2022). Several studies in *D. melanogaster* have provided evidence that sub-lethal infection with DCV increases productivity (Gupta et al. 2017; Kutzer et al. 2023). Within the context of our experiment, the lower reproductive output of the high LT50 lines resulted in low relative fitness, despite their longer survival time. However, under more naturalistic exposure to DCV, if high LT50 lines were investing in immune response, they may be able to clear the infection, resume reproduction, and achieve higher relative fitness.

### Conclusions and future directions

The genetic basis of immune responses and the impact of selection has received considerable theoretical and empirical attention, yet in contrast to other quantitative traits, the input of new variation by mutation is understudied. Our results were partially consistent with a prior study in *C. elegans*, but while the work of Etienne et al. (2015) implicating unconditionally deleterious mutation, our results suggest that mutations contributing to poor survival under infection do not have similarly deleterious effects on reproductive output. Interaction between immune and non-immune fitness is likely to vary among biotic and abiotic environmental conditions that alter the energetic demands on organisms. Deleterious fitness effects of mutations can be masked when resources are abundant (Agrawal and Whitlock 2010). We reared flies under resource-limited conditions shown previously to uncover mutational effects (Kannan et al. 2023). However, we cannot rule out that both the mean LT50 relative to the ancestor, and the relationship between LT50 and productivity may have different patterns under greater resource limitation. Further research is needed to determine whether mutational effects on immune function and fitness are consistent among host and pathogen taxa, and how factors such as sex differences and environment may influence mutational effects.

## Data availability statement

The data underlying this article are available in UQ eSpace at 10.48610/48d7c29

## Author contributions

KM conceived the study with KNJ and BMM. All authors contributed to experimental design; BMM conducted experiments with contributions from AKA and KM. BMM and KM analysed the data with input from AKA and KNJ. BMM and KM wrote the first draft of the manuscript and all authors contributed to the final manuscript.

## Acknowledgments

We thank N. Appleton, D. Sun, L. King, A. Kannan & A. Ichim for generating the MA lines. N. Appleton and D. Chew helped with data collection. E. Hine and V. Narayan provided valuable input to survival analyses.

## Supplementary Information

### Wolbachia screen

*Wolbachia* species of intracellular bacteria are estimated to infect over 60% of insect species (Hilgenboecker et al. 2008), although infection has not been detected in *D. serrata* (Clancy and Hoffmann 1997)*. Wolbachia* influences host traits, including disease resistance (Hedges et al. 2008; Teixeira et al. 2008). To ensure that results of the current study were not influenced by *Wolbachia*, a PCR screen for *Wolbachia* was performed on the ancestor population. *Wolbachia* is vertically (maternally) transmitted (Hoffmann 2020); absence of *Wolbachia* in the ancestor DsGRP line was presumed to indicate MA lines had inherited this *Wolbachia*-free state. We also screened a further four DsGRP lines for *Wolbachia* to assess prevalence in the laboratory population more broadly.

Pools of five flies were homogenised in 200 μl of phosphate-buffered saline (PBS) with three glass beads (Sigma-Aldrich) in a Tissue Lyser II (QIAGEN) at 30 hertz per second for 90 seconds. Samples were then centrifuged at 17 × G for 5 minutes, and the supernatant was transferred to a new tube and mixed with 20 μl of proteinase K (600 milli-absorbance units per millilitre). DNA was extracted from flies using the DNeasy Blood & Tissue Kit (QIAGEN) following the manufacturer’s instructions. *Wolbachia*-infected and tetracycline-treated *Drosophila melanogaster* flies with a w^1118^ background (Bloomington Drosophila Stock Centre, ID: 3605) were used as positive and negative controls for *Wolbachia*, respectively. Tetracycline is an antibiotic used to clear flies of *Wolbachia* and is commonly used to create *Wolbachia*-free lines (Hoffmann et al. 1986).

PCR reactions were set up using the MyTaq DNA Polymerase kit (Bioline), with primers (wsp 81F 5’ TGGTCCAATAAGTGATGAAGAAAC 3’ and wsp 691R 5’ AAAAATTAAACGCTACTCCA 3’) targeting a single copy outer membrane protein gene, *Wolbachia Surface Protein*, WSP, in Wolbachia (Zhou et al. 1998). As a control for DNA quality in each extraction, additional PCR reactions were set up using 18S ribosome primers (q18S-F 5’ CGAAAGTTAGAGGTTCGAAGGCGA 3’ and q18S-R 5’ CCGTGTTGAGTCAAATTAAGCCGC 3’). Reactions were placed in a SimpliAmp Thermal Cycler (ThermoFisher) under the following conditions: 35 cycles of 94°C 1 min, 55°C 1 min, 72°C 1 min. For each DNA sample, the two PCR products (from the WSP and 18S primers) were combined and then visualised on a 2% agarose gel using a 100 bp DNA Ladder (New England Biolabs).

*Wolbachia* was not detected in any of the *D. serrata* DsGRP lines (Figure S1, no WSP band in lanes 1-5), despite good quality DNA being obtained from all samples (18S band in all samples), and the assay successfully detecting *Wolbachia* when it was known to be present (WSP band in *D. melanogaster* positive control, lane 7). We therefore conclude that the MA experiment was free from influence of *Wolbachia* infection.

**Figure S1.**
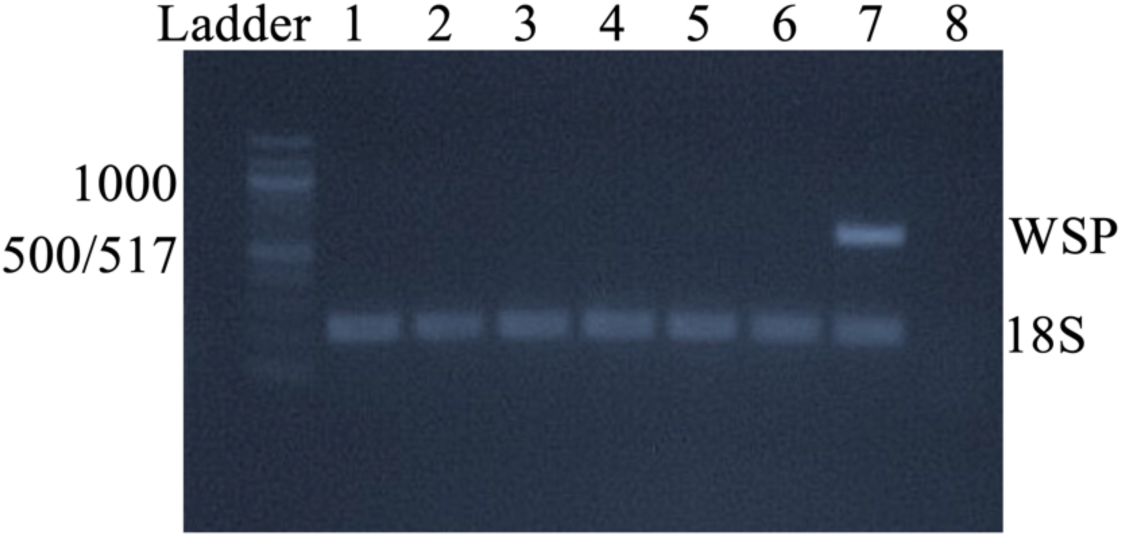
Image of PCR gel from *Wolbachia* detection assay. Separate PCR reactions were run for the WSP and 18S primers, and products were combined before loading onto the gel. Lanes 1 through 5 contained PCR products from DsGRP lines: DsGRP-63, DsGRP-20, DsGRP-120 (the MA ancestor), DsGRP-101 and DsGRP-170, respectively. Lanes 6 and 7 contained PCR products from tetracycline-treated (lane 6) and *Wolbachia*-infected (lane 7) *D. melanogaster* (w^1118^ background), which comprised the negative and positive controls for *Wolbachia*, respectively. Lane 8 contained a further negative control for DNA contamination: water only. A 100bp DNA ladder (New England Biolabs) was used to determine PCR product sizes, and sizes of major ladder bands are displayed to the left of the gel in base pairs (bp). To the right of the gel image, “WSP” indicates the expected position of bands for WSP primers, for which the PCR product is approximately 500 bp, varying slightly among *Wolbachia* strains (Zhou et al. 1998), while “18S” indicates the expected position of 18S PCR products at 200 bp.

### Effect of DCV in males

To determine if males were similarly susceptible to DCV infection, survival assays were conducted on both males and females from the ancestral DsGRP-120 line. These assays were performed simultaneously to the assay of 15 randomly chosen MA lines, following identical procedures (see main Methods Experiment 1). Twenty replicate vials of seven virgin males or seven virgin females (40 vials in total) were injected with PBS or with DCV over two consecutive days, with equal numbers of vials assigned to each treatment and each injection day (five PBS and five DCV vials per sex per day).

We fit a Cox proportional hazards mixed effects model using the *coxme* package (Therneau 2022) in R (R Core Team 2022; Posit Team 2023). Treatment (PBS or DCV injected), sex (male or female), the interaction between treatment and sex, and injection day were modelled as categorical fixed effects, while replicate vial was modelled as a random effect. We used a log-likelihood ratio test (LRT), comparing the fit (-2 log likelihood) of the full model to a nested model in which the interaction between injection treatment and sex was not fit; the difference in fit between the nested models follows a chi-square distribution with degrees of freedom equal to the difference in number of model parameters (here, 1) (Self and Liang 1987). We followed the same approach to apply LRT to each of the main effects.

We accepted the null hypothesis that DCV increased mortality relative to PBS to the same extent in both sexes (LRT of sex by treatment interaction: χ^2^ = 0.72, d.f. = 1, *P* = 0.3975; Figure S2). Survival time in both sexes was negatively affected by DCV injection (LRT of injection treatment: χ^2^ = 225.58, d.f. = 1, *P* < 0.0001; Figure S2). Males, on average, died earlier than females, irrespective of injection treatment (LRT of Sex: χ^2^ = 21.74, d.f. = 1, *P* < 0.0001; Figure S2). The mean LT50 for males and females was 4.72 and 5.94 days, respectively. Earlier male mortality may reflect impacts of physical damage arising via aggressive male interactions within the single-sex vials (K. McGuigan, pers. obs.). A similar sex difference in overall survival, rather than DCV morbidity, has been reported following sublethal infection with DCV in *D. melanogaster* (Gupta et al. 2017).

**Figure S2.**
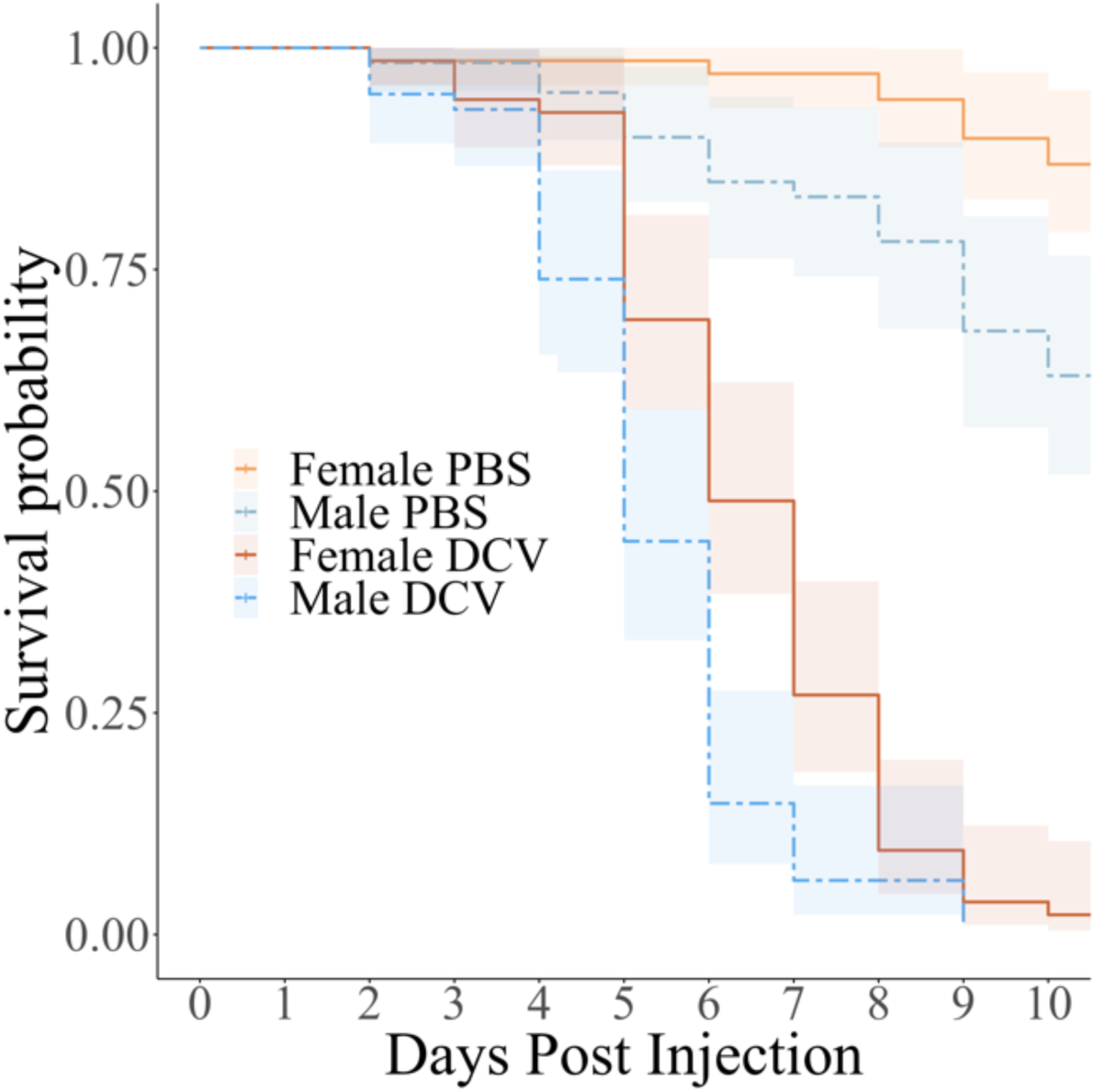
Survival of male and female *D. serrata* DsGRP-120 flies challenged with DCV or PBS. The probability of being alive each day was estimated from a Cox proportional hazard mixed effects model, with treatment (PBS control, lighter coloured lines or DCV pathogen infected, darker coloured lines) and sex (female, solid lines, or male, dotted lines) as the predictors. Shading indicates 95% confidence intervals. All DCV injected flies were dead by day 11, and PBS flies were censored.

### Mean effect of mutation on survival following DCV injection

As detailed in the main Methods and Results, we assessed the mean effect of mutation on survival following DCV injection via comparison of LT50 values between MA and DsGRP-120. Here, we compliment that result with analysis of data from Experiment 1: females from 15 randomly chosen MA lines and DsGRP-120 were injected with either PBS or DCV. We fit a Cox proportional hazards mixed effects model in which treatment (PBS or DCV injected), genotype (DsGRP-120 ancestor or derived Mutation Accumulation, MA), the interaction between treatment and genotype, and injection day were modelled as categorical fixed effects, while line and replicate vial (nested within line and treatment) were both modelled as random effects. To test the null hypothesis of unbiased mutational effects, we compared the fit (-2 log likelihood) of this model and a model in which the treatment by genotype, or genotype effects were not modelled using LRT, as detailed above.

We accepted the null hypothesis that DCV affected mortality to the same extent in both the ancestor and derived MA lines (LRT of genotype by treatment interaction: χ^2^ = 0.28, d.f. = 1, *P* = 0.5937; Figure S3). There was also no evidence of any effect of mutation on the mean survival time, irrespective of injection treatment (LRT of genotype: χ^2^ = 0.45, d.f. = 1, *P* = 0.5013; Figure S3).

**Figure S3.**
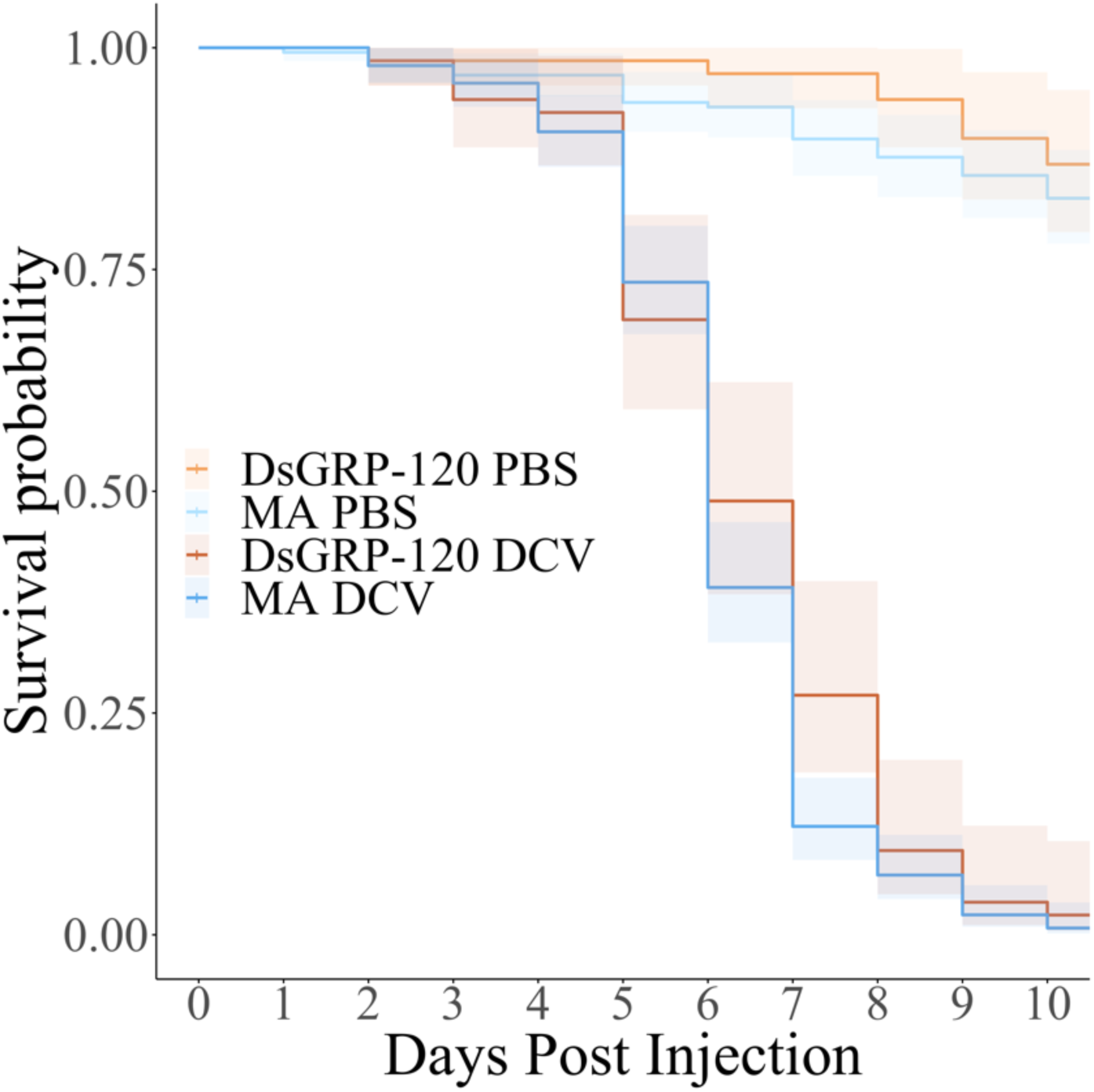
Survival of MA line or DsGRP-120 females challenged with DCV or PBS. The probability of being alive each day was estimated from a Cox proportional hazard mixed effects model, with treatment (control, PBS, lighter coloured lines versus pathogen, DCV, darker lines) and genotype (DsGRP-120 Ancestor or the 15 derived MA lines) as the predictors. Shading indicates 95% confidence intervals. All DCV injected flies were dead by day 11, and PBS flies were censored.

